# Comparative mode of action of antimicrobial peptide melimine and its derivative Mel4 against *Pseudomonas aeruginosa*

**DOI:** 10.1101/450577

**Authors:** Muhammad Yasir, Debarun Dutta, Mark DP Willcox

## Abstract

Melimine and Mel4 are chimeric cationic peptides with broad spectrum antimicrobial activity, and recent investigations have shown that they are highly biocompatible with animal model and human clinical trials. The current study examined the mechanism of action of these two antimicrobial peptides against *P. aeruginosa* with a series of investigations. Antimicrobial activities were determined by MIC and MBC. Endotoxin neutralization was determined using the LAL assay, effect on the cytoplasmic membrane was evaluated using DiSC(3)-5 and Sytox green stains, and Syto-9 and PI dyes using flow cytometry. Release of cytoplasmic materials (ATP and DNA/RNA) were determined using ATP luminescence and increase in OD_260nm_. The ability to lyse bacteria was studied by measuring a decrease in OD_620nm_. The MIC of the peptides remained low against *P. aeruginosa* strains, which showed efficient neutralization of LPS, indicating their role in the anti-pseudomonas and LPS binding activities. Both AMPs rapidly (starting at 30 seconds) depolarized *P. aeruginosa* cytoplasmic membrane leading to reduction in viability. Melimine was responsible for more ATP release (75%) compared to Mel4 (36%) (*P*<0.001) following two minutes exposure. For both peptides, Sytox green entered cells after five minutes of incubation. Flow cytometry demonstrated that both the AMPs permeabilized the cell membrane at 30 minutes and followed by increasing permeability. Similar results were found with DNA/RNA release experiments. Overall, melimine showed higher ability of membrane disruption, cell lysis compared to Mel4 (*P*<0.001). Knowledge regarding mechanism of action of these two AMPs would be helpful in making them as anti-pseudomonas drug.

## INTRODUCTION

*Pseudomonas aeruginosa* is a metabolically versatile ubiquitous Gram-negative opportunistic pathogen that can cause infections in animals and plants (1). *P. aerugino*sa is responsible for localized to systemic infections in humans, which can be life threatening. Over the years, *P. aerugino*sa has become one of the most frequent causative agents of nosocomial infections, associated with substantial morbidity and mortality (2). The current standards of care to treat *P. aeruginosa* infections are not effective (3) as its outer membrane offers low permeability, which limits the penetration of antibiotics into the bacterial cells thereby increase antibiotic resistance (4, 5). Given the severity of *P. aeruginosa* infections and the limited antimicrobial arsenal with which to treat them, finding new alternative antimicrobials with unique mechanisms of action is urgently required (6).

Antimicrobial peptides (AMPs) are part of the innate immune response of living organisms and have broad spectrum activity ranging from viruses to parasites at low concentrations (7). AMPs are usually cationic in nature and have a varying number (from five to over a hundred) of amino acids. AMPs possess multiple modes of action, rapid bacterial killing kinetics and little toxicity toward human cells (8, 9). Bacteria do not easily gain resistance to AMPs due to their fast killing mechanism and multiple target sites (10, 11). The mechanism of action of AMPs starts by interacting with negatively charged lipopolysaccharides (LPS) in the outer membranes of Gram negative bacteria (12-14) leading to a destabilization and permeabilization (15). AMPs are then able to interact with the cytoplasmic membrane. Several models for the interaction of AMPs with bacterial cytoplasmic membranes have been proposed, such as “barrel stave” “toroidal pore” and “carpet model” (12). In all of these models AMPs displace lipids, alter membrane structure and these interactions result in leakage of cellular contents such as K+, ATP and DNA/RNA, ultimately resulting in cell death (16, 17). AMPs can also act intracellularly disrupting DNA, RNA and protein synthesis (18, 19).

Melimine (TLISWIKNKRKQRPRVSRRRRRRGGRRRR) is a cationic hybrid peptide of melittin and protamine (20). It has broad spectrum activity against a wide range of Gram-negative and Gram-positive bacteria (including MRSA and multi drug resistant *P. aeruginosa*, fungi and protozoan such as *Acanthamoeba*) (20, 21). Importantly, bacteria did not develop resistance against melimine when exposed at sub-MIC for 30 consecutive days (20). Moreover, melimine is not cytotoxic at well above active concentrations (20,21). Melimine has a random coil confirmation in aqueous environments but increases its ⍺- helical content (to 35-40%) in bacterial membrane mimetic environments. This change in conformation upon binding to the membrane is widely accepted as the first step in the mechanism of action of many AMPs (22). Melimine disrupts the outer membrane and rapidly destroys the inner membrane potential of *P. aeruginosa* (23). Melimine retains its antimicrobial activity when bound to polymers and titanium (20, 21, 24). However, in a human clinical trials, melimine-coated contact lenses caused corneal staining (25).

A shorter sequence of melimine called Mel4 (KNKRKRRRRRRGGRRRR) has been designed which was shown to not result in corneal staining after being bound to contact lenses (26). Mel4 is highly active against *P. aeruginosa* in solution or when immobilized on surfaces (27). It is non-cytotoxic to mammalian cells *in vitro* (20, 21), animal model studies in vivo and in human clinical trials (26, 28). As can be seen from the amino acid sequence of both peptides, Mel4 has had several amino acids removed, including the single tryptophan in melimine. Tryptophan is known to be a highly lipophilic amino acid (29), and many cationic peptides contain tryptophan as an important part of their mode of action (30-32). Similarly, other non-polar amino acids such as Leu and Ile can encourage peptide binding and disruption of cell membranes (33). Given that both the peptides have been extensively investigated with human clinical trials, their bactericidal mechanism is relatively unknown. In addition, amino acid sequence of Mel4 is different to melimine, its mechanism to kill Gram negative bacteria is completely unknown. Hence, this study examined and compared the mode of action of melimine and Mel4 against *P. aeruginosa*.

## MATERIALS AND METHODS

All experiments were run in triplicate and repeated on three separate occasions, except for flow cytometry data which were obtained after two repeats. For all experiments, negative controls for the effect of melimine and Mel4 were simply bacterial cells grown in their absence.

### Synthesis of peptides

Melimine and Mel4 (≥90% purity) used in the current study were synthesized by conventional solid-phase peptide protocols and procured from the Auspep Peptide Company (Tullamarine, Victoria, Australia).

### Bacterial strains

Different strains of *P. aeruginosa* such as 6294 and 6206 (microbial keratitis isolates; 6294 an invasive strain containing the *exoS* gene and 6206 a cytotoxic strain containing the *exoU* gene) (34), Paer1 (isolated from contact lens induced acute red eye, contains the *exoS* gene but does not manifest the associated invasive phenotype) (34) and ATCC 19660 (isolated from human septicaemia; a cytotoxic strain containing the *exoU* gene) (35) were used in the current study. All these strains were obtained from stock cultures preserved at −80 °C in Brain Heart Infusion (Oxoid, Basingstoke, UK) containing 25% glycerol.

### Bacterial cell preparation

Bacteria were grown in Tryptic Soy Broth (TSB; Oxoid) for 12-16 h and cells were then washed with phosphate buffer saline (PBS, NaCl 8 g/L, KCl 0.2 g/L, Na_2_HPO_4_ 1.4 g/L, KH_2_PO_4_ 0.24 g/L) and diluted into the same buffer containing 1/1000 TSB to OD_600nm_ 0.05- 0.06 (1× 10^7^ colony forming units (CFU)/ml confirmed upon retrospective plate counts on TS agar (Oxoid). This inoculum preparation was used in most experiments except for assessing the minimum inhibitory and bactericidal concentrations and measuring the release of DiSC3-5 from cells (which was performed in cells (1× 10^7^ CFU/ml) suspended in HEPES buffer).

### Minimum inhibitory and bactericidal concentrations

The minimum inhibitory and minimum bactericidal concentrations of melimine and Mel4 were determined for all strains using a modified version of the Clinical Laboratory and Standard Institute (CLSI) broth microdilution method as reported previously (36), using Muller Hinton Broth (Oxoid) containing 0.01% v/v acetic acid (Sigma Aldrich, St Louis, MO, USA) and 0.2% w/v bovine serum albumin (Sigma Aldrich; MHB). Bacterial cells, diluted to 5×10^5^ CFU ml^−1^ in MHB, were incubated with various concentrations of the peptides. The MIC was set as the lowest concentration of peptides that reduced bacterial numbers by ≥ 90% while the MBC was set as the lowest concentration of peptides that reduced bacterial growth by >99.99% after enumeration of viable bacteria by plate count method compared to bacteria grown with no antimicrobial agent.

### Interaction with *P. aeruginosa* lipopolysaccharide

A limulus amoebocyte lysate (LAL) assay was performed to determine the interaction of AMPs with lipopolysaccharides (LPS) of *P. aeruginosa* using a chromogenic assay kit (Cape Cod, E. Flamouth, MA, USA). Briefly, 8×10^−4^ nmol/ml LPS from *P. aeruginosa* 10 (Sigma Aldrich, St Louis, MO, USA) was dissolved with melimine and Mel4 at 1x or 2x MIC in endotoxin free water (Sigma Aldrich, St Louis, MO, USA) and incubated at 37 ^o^C for 30 min. The interaction of LPS with melimine or Mel4 was assessed as the decrease in OD_405nm_, following addition of the LAL reagent, compared with control (LPS in endotoxin free water) without peptides.

### Cytoplasmic membrane disruption

Three assays were performed to determine whether melimine and Mel4 could affect the cytoplasmic membrane of *P. aeruginosa*. The DiSC3-5 assay was used to determine the effect of the peptides on membrane potential. Two assays, Sytox Green and Propidium Iodide, were conducted to determine whether the peptides could damage cytoplasmic membranes and allow the stains to penetrate and bind to intracellular nucleic acids. Sytox Green has a molecular mass of 213.954 g/mol and a topological polar surface area of 28.7 A^2^, whereas Propidium Iodide has a molecular mass of 668.087 g/mol and a topological polar surface area of 55.9 A^2^(37). Differences in the penetration of these two dyes through the cytoplasmic membranes may be associated with different sizes of pores formed by the AMPs.

Cytoplasmic membrane depolarization by the AMPs was performed as described previously (23) with melimine and Mel4 at 1x and 2x MIC at the final concentrations. The number of viable cells were confirmed by serially diluting aliquots of bacteria in D/E neutralizing broth (Remel, Lenexa, KS, USA) and plating these onto Tryptic Soy Agar (Oxoid, Basingstoke, UK) containing phosphatidylcholine (0.7 g /L) and Tween 80 (5ml/L). The plates were incubated at 37 ^o^C overnight and the number of live bacteria were enumerated and expressed as CFU/ml. Two positive controls of dimethyl sulfoxide (DMSO) (Merck, Billerica, MA, USA) (20% v/v) in HEPES (100 µl) and sodium azide were used to depolarise the cytoplasmic membranes of bacteria (38).

For Sytox green penetration into cells, the protocol was adopted from Li *et al.,*(39) with a few modifications. Briefly, bacterial cells, aliquots (100 µl) were dispensed into wells of 96-well plates along with 5µM Sytox green (Invitrogen, Eugene, Oregon, USA). Plates were incubated for 15 minutes in the dark at room temperature and then 100 µl of melimine and Mel4 were added equal to 1x or 2x as final concentrations. The increase in fluorescence was measured spectrophotometrically (at an excitation wavelength of 480 nm and an emission wavelength of 522 nm) every 1 minutes up to 30 minutes, and then after 150 minutes. A positive control of 1% (v/v) Triton X-100 (Sigma Aldrich, St Louis, MO, USA) in PBS with 1/1000 TSB (100 µl) was used to disrupt the cytoplasmic membrane of bacteria.

Flow cytometry was used to quantify the ability of melimine and Mel4 to permeabilize bacterial membranes of *P. aeruginosa* 6294 resulting in incorporation of propidium iodide (PI) (Invitrogen, Eugene, Oregon, USA) into cells with compromised cell membrane. Bacterial cells were stained simultaneously with SYTO9 and PI at concentrations of 7.5 µM and 30 µM respectively and incubated at room temperature for 15 min. Fluorescence intensities were recorded with LSRFortessa SORP Flow cytometer after addition of 1x and 2x MIC of melimine or Mel4 at different time points. The wavelength of green fluorescence was (525/550 nm) bandpass filter for SYTO9 and a red fluorescence (610/20 nm) bandpass filter for PI (40). Data were acquired and analyzed using Flowjo software (USA). Minimum 20000 events were recorded for each sample.

### Leakage of intracellular contents

The leakage of ATP and DNA/RNA was measured in separate assays. Aliquots of 100 µl of bacteria were incubated with melimine at the final concentrations equal to 1x or 2xMIC and at 37°C for 10 min. The samples were taken at 2 min intervals and centrifuged at 9000 × g for five minutes and the supernatant was kept on ice until further use. For determination of internal ATP the bacterial pellets were resuspended in boiling 100 mM Tris, 4 mM EDTA pH (7.4) and further incubated for 2 mins at 100 °C to lyse all the cells. The lysed cells were centrifuged at 9600 × g for two minutes and the supernatant was kept on ice until further analysis (41). Subsequently, both total and extracellular ATP were determined using an ATP bioluminescence kit (Invitrogen, Eugene, Oregon, USA) according to manufacturer’s instructions.

The assay for measuring the loss of DNA/RNA was performed according to protocol Carson *et al.,* (42) with some modification. Aliquotes (100 µl) of bacteria was incubated with melimine and Mel4 at their 1x or 2x MIC and incubated at 37 ^o^C. Samples were collected at different time intervals, diluted (1:10) and filtered through 0.22 µm pores (Merck, Tullagreen, Ireland). The OD_260nm_ of the filtrates was measured in UV-star plate (Greiner Bio-one GmbH, Frickenhausen, Germany). The results were expressed as the ratio to the initial OD_260nm_.

### Lysis of bacteria

This experiment was adopted from the method of Carson *et al.,* (42). The bacterial lytic potential of the two peptides was evaluated using two different bacterial inoculums 1× 10^8^ CFU/ml and 3 × 10^10^ CFU/ml. The smaller inoculum size was tested to see whether OD_620nm_ was measurable for 1× 10^8^ CFU/ml. The OD_660nm_ of bacterial suspension was adjusted to 0.1 to yield 1× 10^8^ CFU/ml. The larger inoculum size of 3 × 10^10^ CFU/ml was obtained by adjusting OD_620nm_ 0.3. The bacterial numbers CFU/ml were confirmed by retrospective plate count. Melimine and Mel4 were added at 1x MIC and 2x MIC as final concentrations. Bacterial cultures were immediately mixed and then diluted 1:1000 in TSB. The OD_620nm_ was measured and additional readings were taken at 30, 60, 90, 120 minutes, 6.5 and 24 h after incubating at room temperature. PBS with peptides at their respective concentrations was used as blank. The results were recorded as a ratio of OD_620nm_ at each time point compared to the OD _620nm_ at 0 minutes (in percentage).

### Statistical analyses

Statistical analyses were performed using GraphPad Prism 7.02 software (GraphPad Software, La Jolla, CA, USA). The effect of the different concentrations of peptides was analysed using Tukey’s test of multiple comparisons. Correlations between release of extracellular ATP and bacterial death were examined using Pearson correlation test. Statistical significance was set as *P*<0.05.

## RESULTS

### Inhibitory concentrations of peptides

MICs and MBCs for melimine and Mel4 against *P. aeruginosa* are shown in **Table 1.** Mel4 had highest bactericidal activity against *P. aeruginosa* strains 6294, 6206 and ATCC 19660 with a MIC of 26.6 nmol/ml. The MIC was lower for melimine against strains 6294, 6206 and Paer1 at 66 nmol/ml compared to strain ATCC19660. For Mel4, the MIC was lowest against stains 6206, 6294 and ATCC19660 at 26.6 nmol/ml. For all strains the MBC was usually 2x the MIC except for strain 6294 where the MBC for both melimine and Mel4 was ≥4X the MIC while for ATCC 19660 the MBC for Mel4 was equivalent to the MIC. Melimine at its lowest MBC needed 1.59×10^12^ molecules per cell to cause death whereas Mel4 needed 3.2×10^11^ molecules per cell to cause cell death at its lowest MBC.

**Table 1.** MIC and MBC values of melimine and Mel4 against strains of *P. aeruginosa*

| Bacterial strains | Melimine |  | Mel4 |  |
| --- | --- | --- | --- | --- |
|  | MIC nmolml<sup>-1</sup> (µgml<sup>-1</sup>) | MBC nmolml<sup>-1</sup> (µgml<sup>-1</sup>) | MIC nmolml<sup>-1</sup> (µgml<sup>-1</sup>) | MBC nmolml<sup>-1</sup> (µgml<sup>-1</sup>) |
| *P. aeruginosa* 6206 | 66 (250) | 132 (500) | 26.6 (62.5) | 53.2 (125) |
| *P. aeruginosa* 6294 | 66 (250) | 528 (2000) | 26.6 (62.5) | 106.5 (250) |
| *P. aeruginosa* Paer1 | 66 (250) | 132 (500) | 106.5 (250) | 213 (500) |
| *P. aeruginosa* ATCC 19660 | 132 (500) | 264 (1000) | 26.6 (62.5) | 26.2 (62.5) |

MIC= minimum inhibitory concentration that inhibits the growth of ≥90% of cells
MBC = minimum bactericidal concentration that kills ≥99.9% of cells.

### Interaction with Lipopolysaccharides

Both melimine and Mel4 interacted with the LPS of *P. aeruginosa* and neutralized its endotoxin activity in a concentration dependent manner. At the 66 nmol/ml concentration melimine significantly neutralized LPS which was evidenced by reduction of OD_405nm_ by 1/6^th^ compared to the controls. Similarly, melimine at the concentration of 132 nmol/ml neutralized LPS by 1/8^th^ compared to the controls. Neutralization of LPS was similar with the Mel4 which reduced OD_405nm_ by half (1/2) at 26.6 nmol/ml and 1/4^th^ at 53.2 nmol/ml compare to control.

### Membrane disruption

Both melimine and Mel4 depolarized the cytoplasmic membrane of *P. aeruginosa* in a concentration dependent manner. An increase in fluorescence intensity was detected as early as 30 seconds after addition of either peptide to all the strains of *P. aeruginosa*. **Fig. 1a** shows the data for *P. aeruginosa* 6294, whereas data for all other strains are available in the supplementary documents. Following 150 seconds of exposure, no further increase in the release of DiSC3-(5) was seen for any of the strains investigated. Fluorescein intensity was higher at the MIC of melimine (66 nmol/ml) compared to the MIC of Mel4 (26.6 nmol/ml), and the same trend was observed throughout the time course (*P*<0.005). This depolarization of the cytoplasmic membrane was associated with 3.6 log_10_ and 4.0 log_10_ viable *P. aeruginosa* 6294 inhibition by melimine MIC (66 nmol/ml) and MBC (132 nmol/ml) respectively. Similarly, there was 2.7 log_10_ and 3.2 log_10_ reduction by Mel4 MIC (26.6 nmol/ml) and MBC (53.2 nmol/ml) (**Fig.1b**). Sodium azide also depolarized the cell membrane and released DiSC3-(5) over the period of testing, but with reduced fluorescence intensity and without bacterial killing. Sytox green fluorescence increased over time and was detected as early as 5 minutes (**Fig. 2**). The intensity of fluorescence gradually increased over 30 minutes for all the peptide concentrations for all strains. However, this effect was not concentration dependent. For both melimine and Mel4 no significant differences were observed between the MIC and MBC. At their respective MICs, melimine allowed more Sytox green to enter cells than Mel4 after 150 mins (*P*<0.001). Treatment with the positive control Triton-X 100 (1%) resulted emission of higher Sytox green fluorescence.

**Fig. 1.**
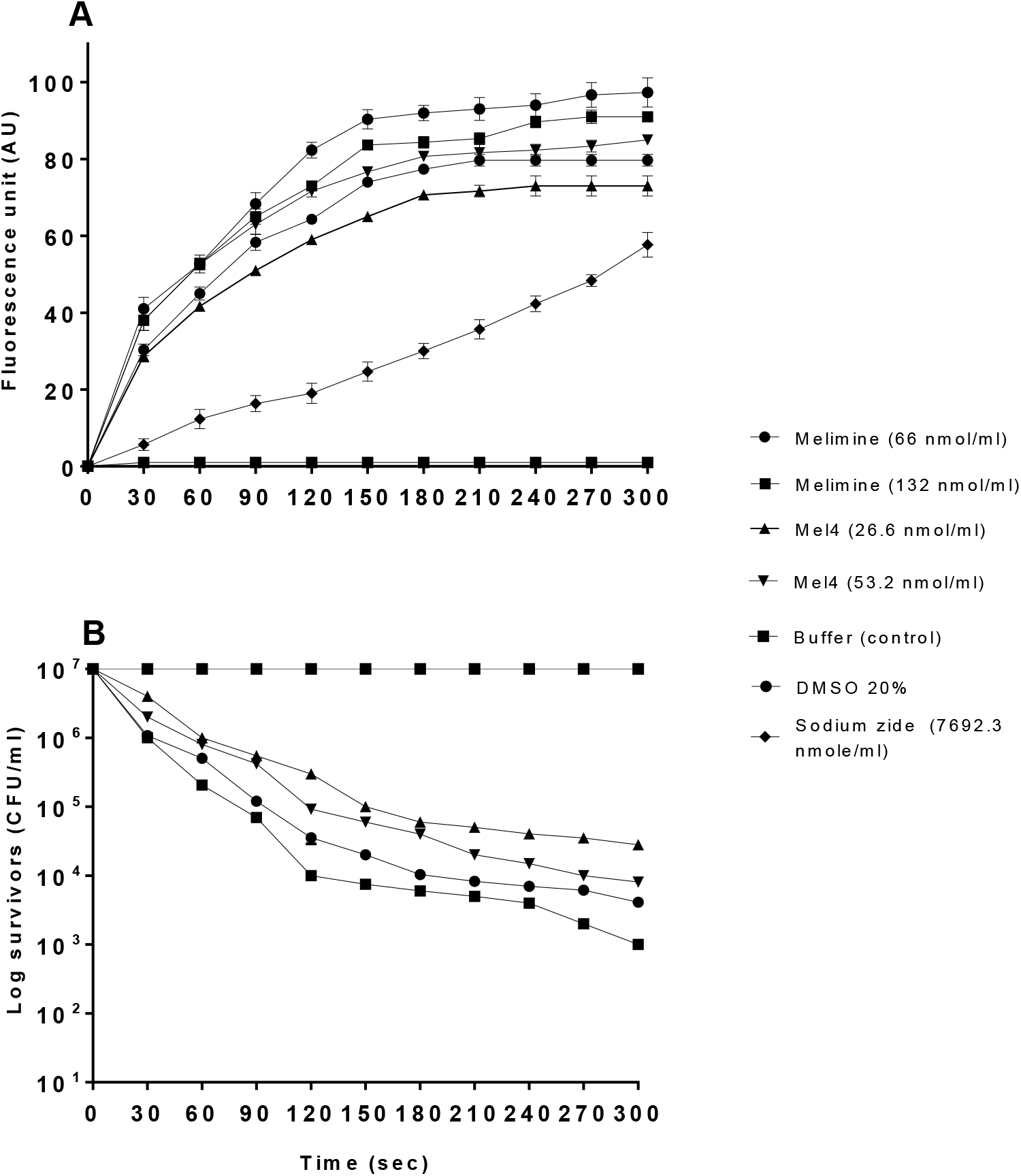
Cytoplasmic membrane depolarization of *P. aeruginosa* 6294 by melimine and Mel4, as assessed by release of the membrane potential-sensitive dye DiSC3-(5) measured spectroscopically at 622_nm_ to 670_nm_ excitation and emission wavelength, and corresponding bacterial survival as determined by plate counts. Data presented as means (±SD) of three independent repeats in triplicate cells. NB, addition of sodium azide to cells did not result in any cell death compared to controls.

**Fig. 2.**
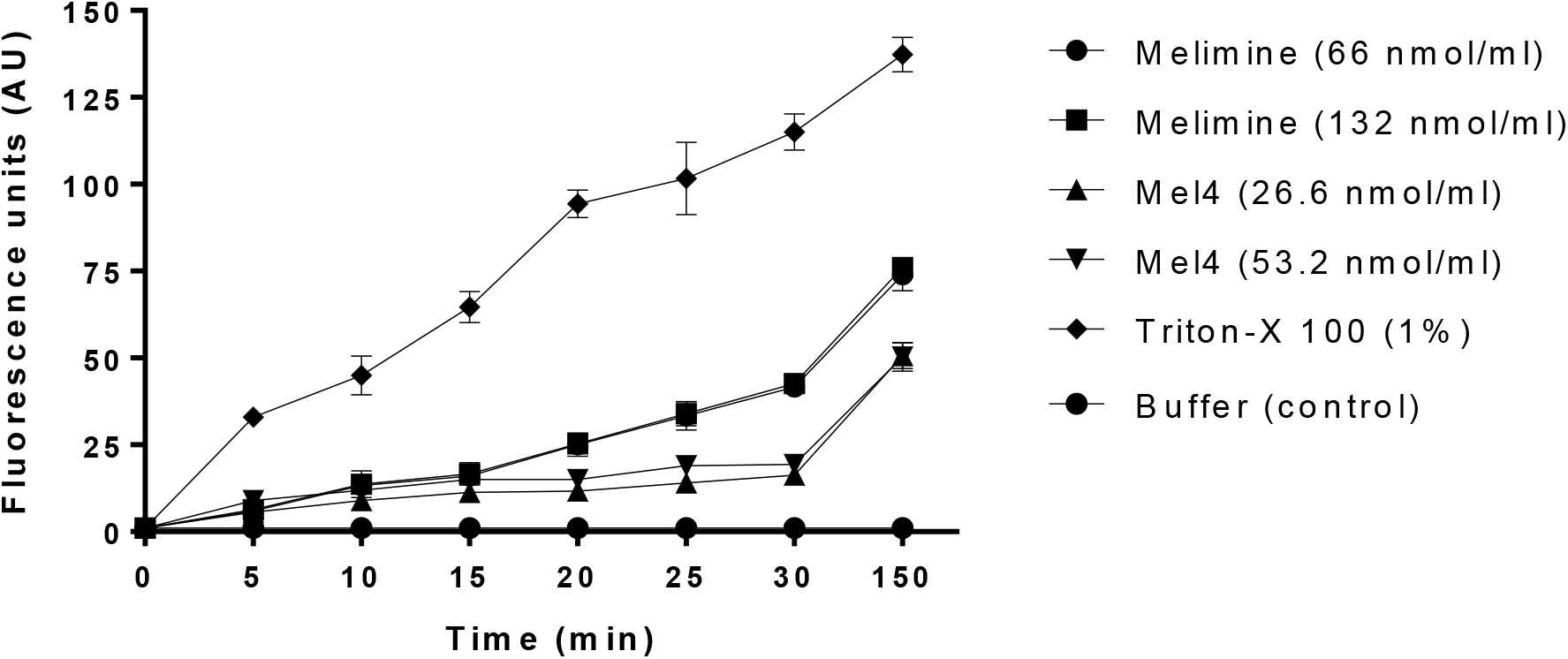
Cytoplasmic membrane permeability of *P. aeruginosa* 6294 by melimine and Mel4 at different concentrations. Fluorescence due to binding of Sytox green fluorescent probe with DNA was measured spectroscopically at 480_nm_ to 522_nm_ excitation and emission wavelength. Data presented as means (±SD) of three independent repeats in triplicate.

The membrane damaging effect of these peptides was also assessed with *P. aeruginosa* 6294 by the flow cytometry in the presence of the DNA intercalating dye PI. The peptides were able to permeabilize the cell membrane in a concentration dependent manner, and this effect was time-dependent with a higher influx of PI after 150 mins incubation (**Fig. 3**). Melimine permeabilized 52.5% of *P*. *aeruginosa* 6294 cells at its MIC at 30 min. after 150 minutes of exposure, PI stained up to 90% of the cells. Although Mel4 permeabilized *P. aeruginosa* membranes, permeabilization at its MIC was less compared with melimine; Mel4 damaged only 17.7% cells after 30 min which increased to 46.7% after 150 min (**Fig.3**). The positive control Triton-X 100 (1%) permeabilized the membrane in 18.7% cells initially which increased up to 34.1% after 130 min. There was also different kinetics of cell death as it can interact with both lipids as well as proteins, create pores and/or remove both lipids/proteins from the membrane. with Triton-X 100. The negative control (buffer-treated) behaved as expected and showed only < 1% PI-stained cells (**Fig.3**).

**Fig. 3.**
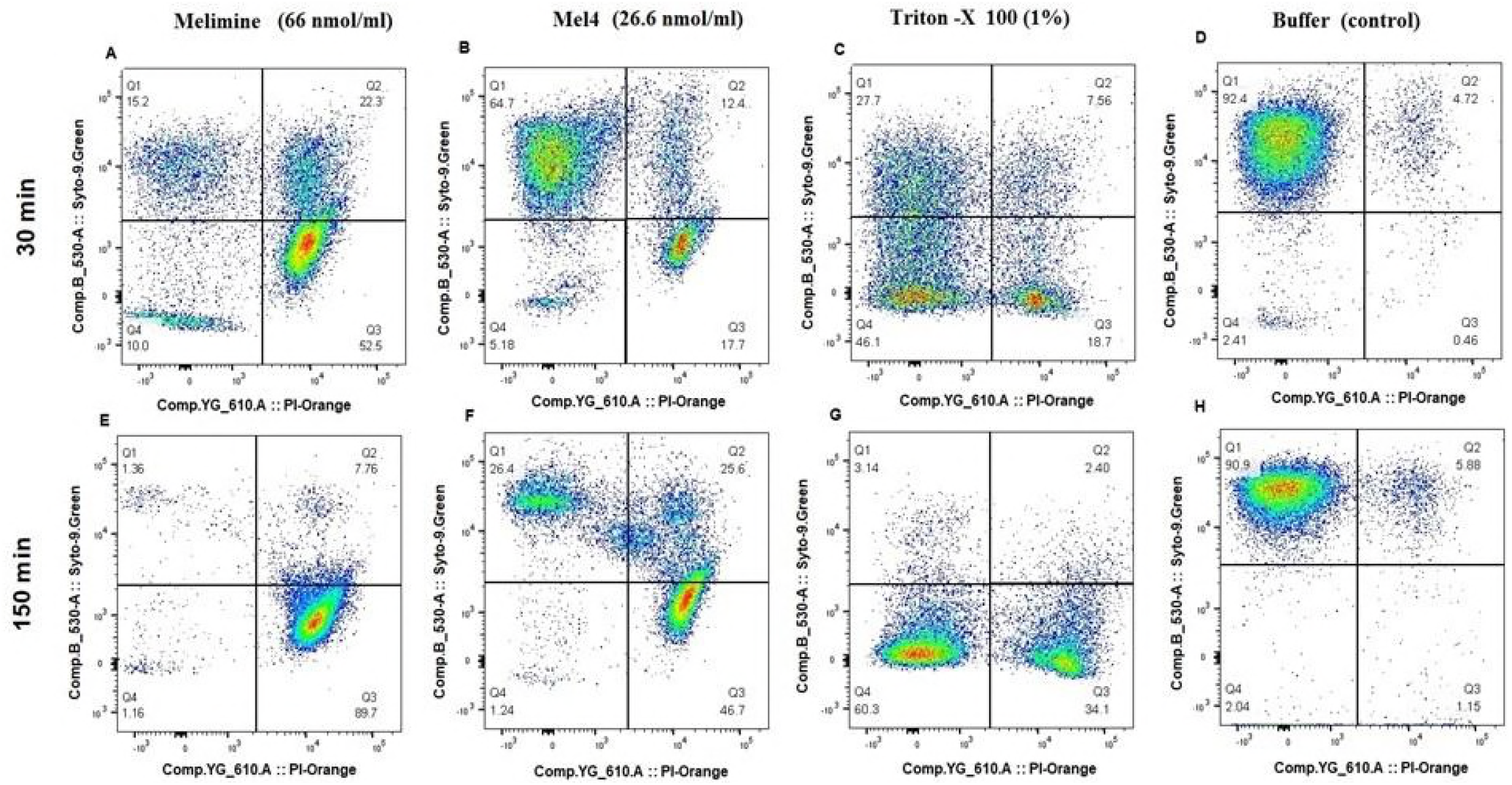
Membrane permeabilization of *P. aeruginosa* 6294 produced by melimine and Mel4 at their MICs determined by flow cytometry with Syto 9 (membrane permeable) and Propidium Iodide (membrane impermeable) stains.

### The release of cytoplasmic contents

Melimine caused 75% and 92 % of ATP release at 1x and 2x MIC concentrations after 2 minutes (*P*<0.001) (Figure 4). The increase of extracellular ATP directly correlated with the loss of viability of *P. aeruginosa* (*R*^2^>0.987). Within the first two minutes, melimine decreased viability by > 3.0 log_10_. Whereas Mel4 released 36% and 44% extracellular ATP at 1x or 2x MIC respectively at 2 minutes (**Fig. 4**). Further incubation for 10 minutes resulted in a slight increase in the release of extracellular ATP. The release of ATP was associated with the reduction of >2.0 log_10_ viable bacteria. Melimine induced more leakage of ATP than Mel4 (*P*<0.001).

**Fig. 4.**
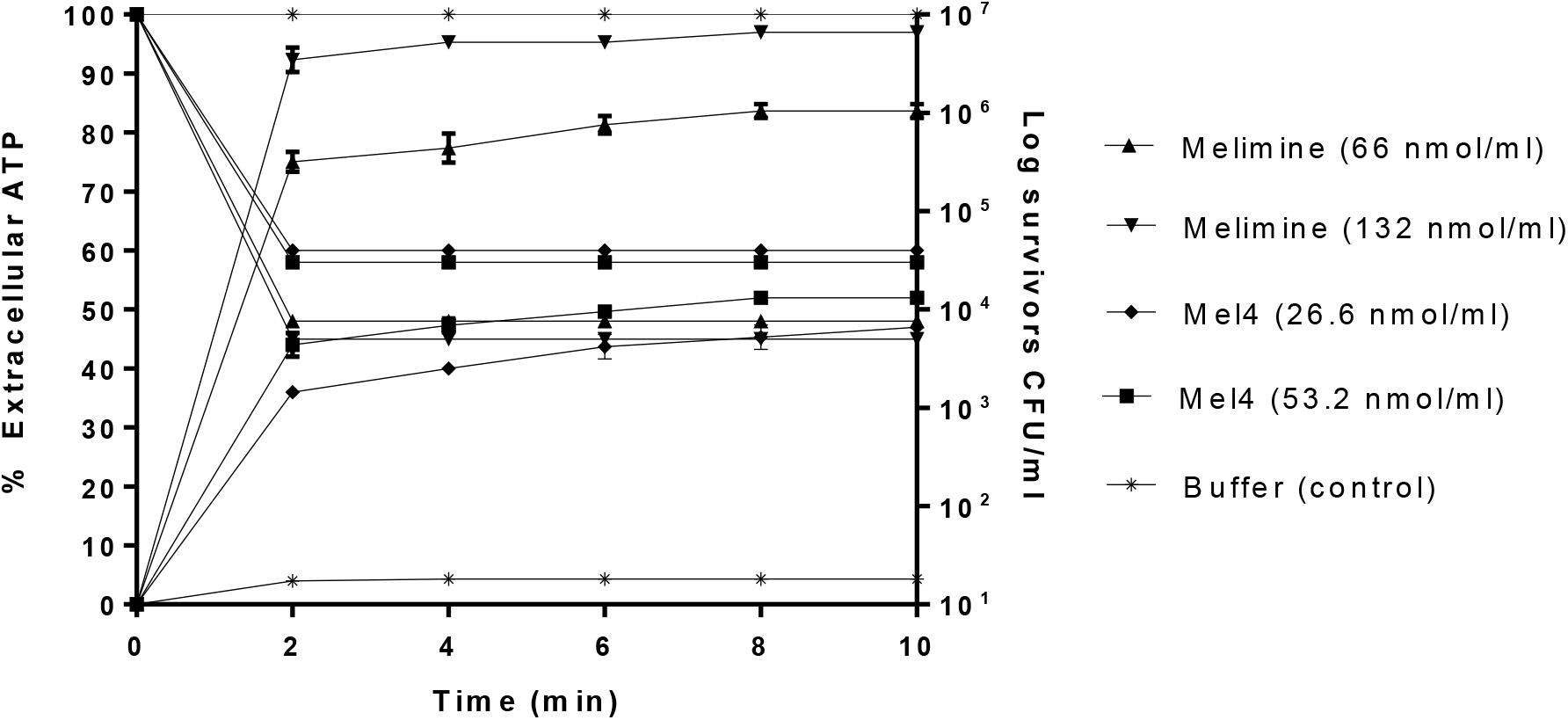
The effect of melimine and Mel4 on ATP release from *P. aeruginosa* 6294 at MIC and two times MIC concentrations and the corresponding change in the number of viable cells. Data presented as means (±SD) of three independent repeats in triplicate compared with buffer-treated control.

**Fig. 5** shows the releases of DNA or RNA (260nm absorbing material) following incubation with the peptides. The release of DNA or RNA of *P. aeruginosa* was first detected at 2 minutes time. Melimine was associated with a dose-dependent release of DNA/RNA which was significantly higher than observed with the control after 5 min (*P*<0.001). On the other hand, the release of DNA/RNA was not affected by the concentration of Mel4. Compared with Mel4, melimine released higher amounts of DNA/RNA at all the time points observed (*P*<0.001).

**Fig. 5.**
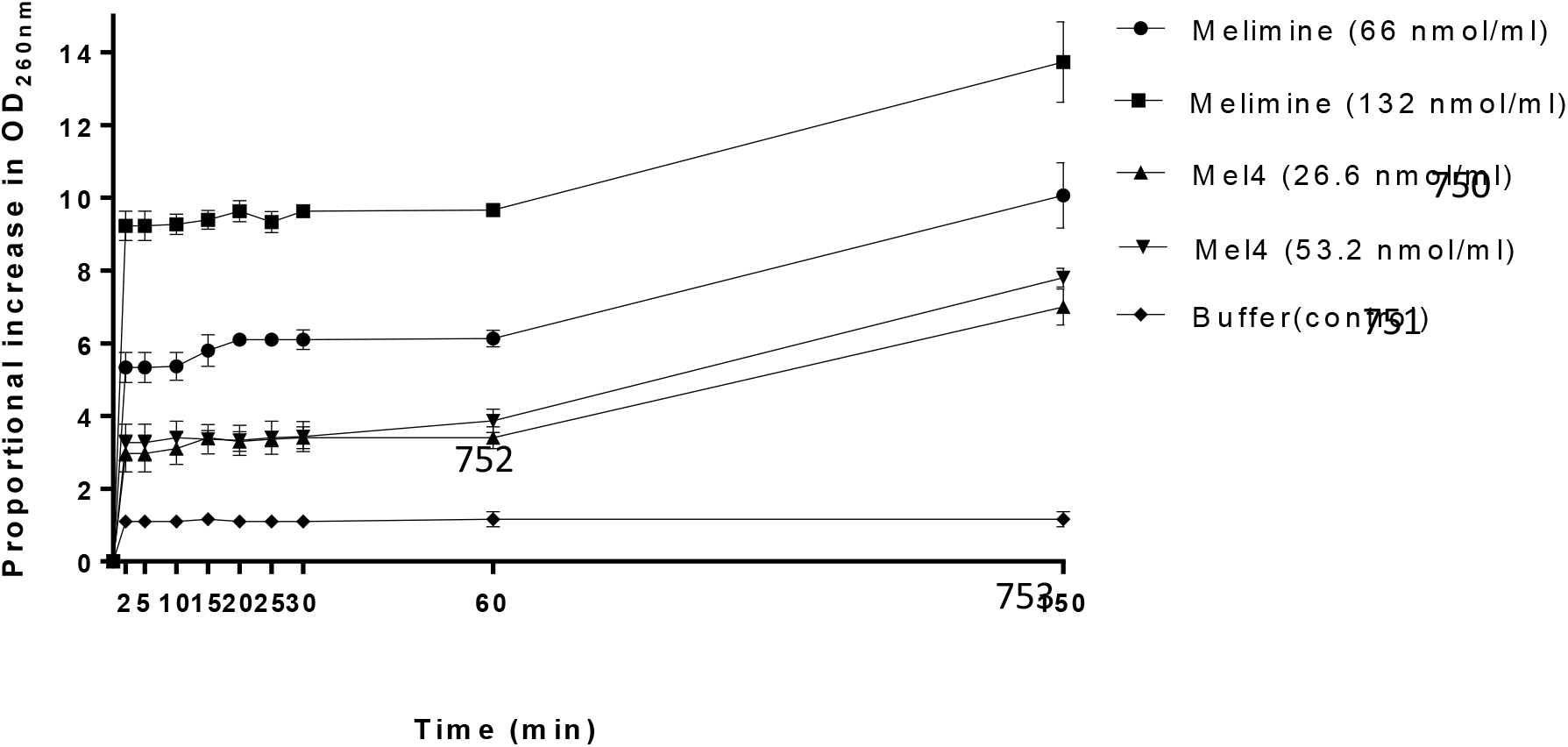
Increase of DNA/RNA from *P. aeruginosa* 6294 due to action of melimine and Mel4 at MIC and two times MIC concentrations determined spectroscopically at OD_260nm._ Data presented as means (±SD) of three independent repeats in triplicate compared with buffer-treated control.

### Bacterial lysis

When the lower cell concentration (1× 10^8^ CFU/ml) was used, no decrease in OD_620nm_ could be seen as the optical density was very low to start with. When higher bacterial inoculum 1× 10^10^ CFU/ml was used and treated with varying concentrations of peptides over 2 hours, a significant reduction in OD_620nm_ was observed (**Fig. 6**). After incubation with melimine, the OD_620nm_ reduced by 25±12% at 1x and 31±3 % at 2x MIC compared to buffer treated negative control (*P*<0.001). OD_620nm_ was further reduced by 37±03% and 52±10% with 1xand 2x MIC after 6 hours respectively. This trend continued over the 24 hours assay where melimine had a higher bacterial-lytic effect and decreased the OD_620nm_ to more than 55±5% at both the concentrations. A similar trend was seen for Mel4 which reduced the OD_620nm_ by 13±8 % and 21±6 % at 1x and 2x MIC respectively after 2 hours (**Fig. 6**). Further reductions in OD_620nm_ by 20±3% with 1x and 48±5% with 2x MIC was observed for Mel4 after 6.5 h respectively. Similarly, OD_620nm_ decreased by 52±2% by both the concentrations of Mel4 after 24 h of incubation (Fig. 6). The bacteriolytic efficiency of both melimine and Mel4 was similar at their MICs (*P*=0.927) but at 2x MIC melimine reduced higher OD_620nm_ than 1x MIC of Mel4 (*P*<0.004) after 24 hours of incubation. The OD_620nm_ of control cells without any peptides remained unchanged over the 24 hours of the experiment.

**Fig. 6.**
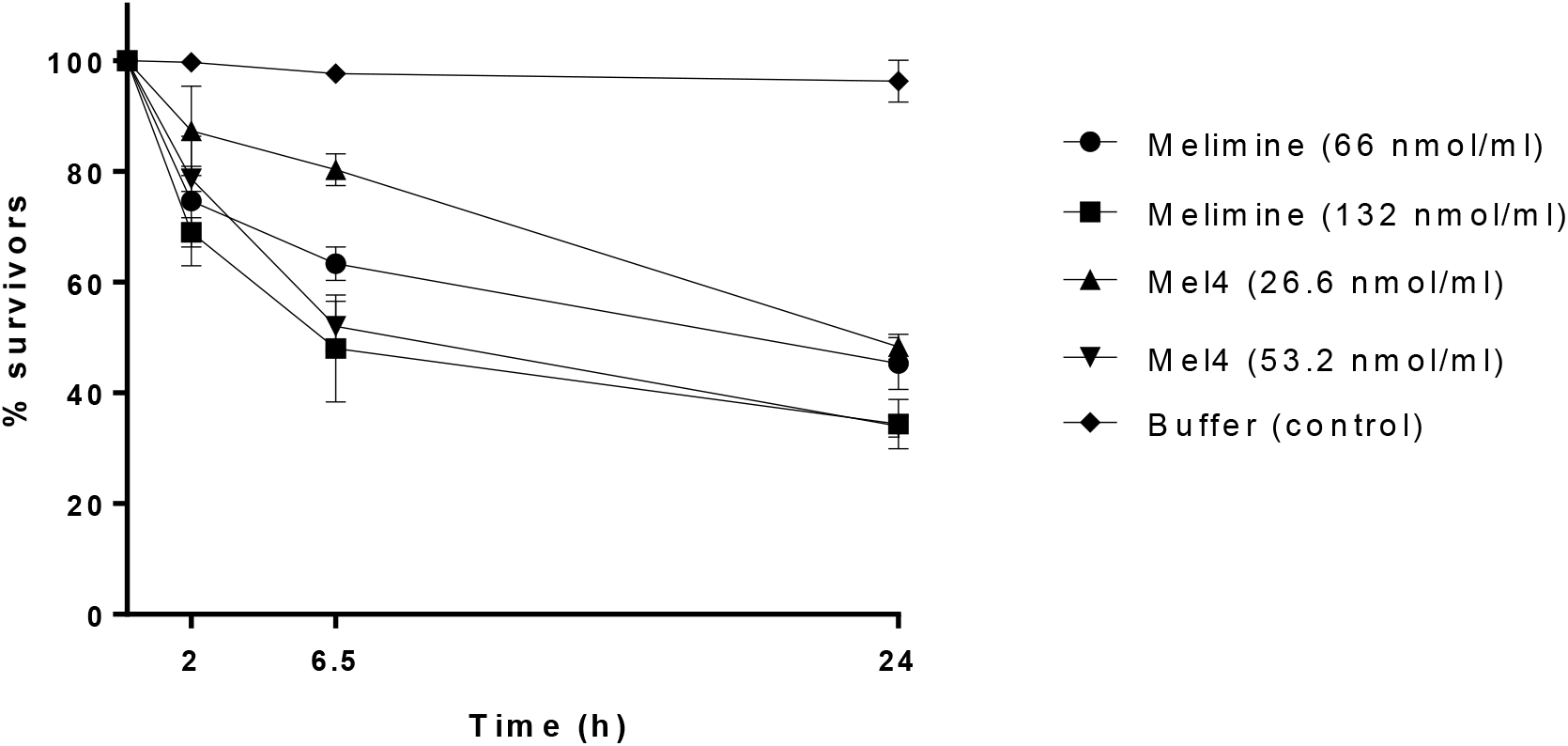
Lysis of *P. aeruginosa* 6294 by melimine and Mel4 at MIC and two times MIC measured spectroscopically at OD_620nm._ Data presented as means (±SD) of three independent repeats in triplicate compared with buffer-treated control.

**Fig. 7.**
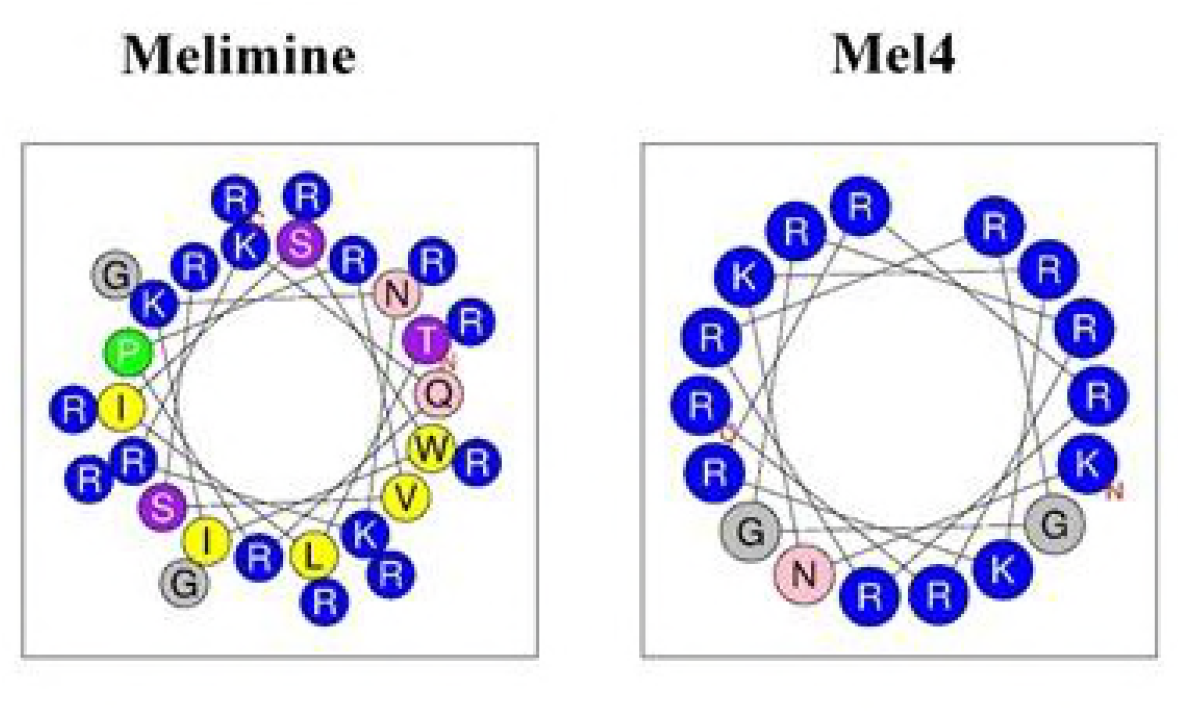
Helical wheel projection of melimine and Mel4. Positive charged residues are represented in blue circles, uncharged residues are in grey circles, polar residues in pink circles and hydrophobic residues in yellow circles.

## DISCUSSION

Over last decade, melimine and Mel4 have been extensively studied *in vitro*, with animal models and human clinical trials. This study for the first time compared and contrasted the timeline of the antimicrobial mechanism of actions of these two thoroughly investigated cationic peptides. The following **Fig. 8** summarizes the sequences of membrane interactions and bactericidal events of the two peptides against *P. aeruginosa* 6294 over the first 2 hours (120 minutes) of exposure. The effects of each of the peptides on all the strains of *P. aeruginosa* were similar. Both peptides were able to significantly neutralise the endotoxic activity of LPS. Both peptides could depolarize the cytoplasmic membrane, and this was associated with rapid loss of cell viability. It was found that loss of viability was quicker with melimine than Mel4. Cytoplasmic membrane depolarisation was followed by ATP and DNA/RNA release from cells, which in turn was followed by permeabilization of the membrane to Sytox green dye that binds to intracellular DNA. When incubated with peptides for 24 hours 50-60% of cells were lysed.

**Fig. 8.** Timeline of bacterial killing by melimine and Mel4. Both AMPs started cell membrane depolarization at 30 seconds followed by ATP and DNA/RNA release at 2 minutes. Cell membrane permeabilization happened at 5 minutes of exposure. Complete bacterial lysis started at 120 minutes of incubation with peptides. All the events started at same time point for both AMPs but intensity of event to occur for melimine was higher than Mel4 at each studied time point.

We have found that melimine has higher bactericidal efficacy when compared to Mel4 at the studied timepoints, and this may be due to their difference in size and structure. The amino acid length required for the peptides to span bacterial cytoplasmic membranes is approximately 15–20 residues, which may vary depending on the thickness of the lipid bilayer (43-45). Mel4 has 17 amino acid, just within the membrane spanning length, whereas melimine has 29 amino acid which probably allows it to easily span the cytoplasmic membrane of *P. aeruginosa*. The shorter length of the Mel4 peptide may mean that it takes longer to penetrate through the outer membrane or start to interact with the inner membrane of *P. aeruginosa* in order to kill the bacteria, or it needs to orientate itself more effectively into the membrane to begin to exert its affects.

Another possible reason behind the differences in the activity of the two peptides is the presence of tryptophan in the sequence of melimine. Tryptophan (Trp) is known to interface with lipid bilayers and can enhance peptide-membrane interactions and facilitate insertion into the membrane (46, 47). A helical peptide RW-BP100 possess a Trp residue which confers higher affinity and deeper insertion into bacterial membranes (48). In addition, Trp has been shown to facilitate the insertion of arginine into the hydrophobic region of membranes *via* cation–π interactions causing rapid membrane disruption (30). Melimine, partly due to the presence of Trp, adopts a partial α helix in bio-membrane mimetic environment (23). A higher helical conformation of peptides is better suited for their antimicrobial activity (49). As Mel4 lacks tryptophan in its sequence, it may have less affinity towards phospholipid bilayers (50). Furthermore, the amino acid sequence of Mel4 predicts that it would have a very low hydrophobic moment (0.039; Table 2) meaning that it is less likely to be attracted within lipid bilayers (51, 52). Also, helical wheel projections of melimine and Mel4 show that the hydrophobic amino acids in melimine segregate to one side of the molecule, whereas as there are no hydrophobic amino acids in Mel4 that can segregate (Figure 7). The lack of the non-polar amino acids Ile and Leu which can encourage peptide binding and disruption of cell membranes (33) may also affect the initial mode of action of Mel4. A recent study (53) has demonstrated that Mel4 does not interact with lipid spheroids composed of 1-oleoyl-2-hydroxy-sn-glycero-3-phosphocholine (PC 18:1) or tethered lipid bilayers composed of 70% zwitterionic C20 diphytanyl-glycero-phosphatidylcholine lipid and 30% C20 diphytanyl-diglyceride ether whilst melimine can interact with these lipid layers. The evidence from the current study suggests that it takes more time for Mel4 to interact with the bacterial membranes to the same extent as melimine due to the changes in its amino acid sequence and subsequent changes in its structure. Indeed, for lipid depolarisation at their respective MICs, it took Mel4 approximately 30 seconds longer to achieve the same degree of depolarisation and 60 seconds longer to achieve the same degree of killing with Mel4 compared to melimine as, evidences by the comparative timeline (**Fig. 1a & 1b**). To permeabilize the cytoplasmic membrane to allow Sytox green to enter cells, Mel4 at its MIC took more than 30 minutes to achieve the same degree of permeabilization that melimine did at its MIC at 5 min. Confirmation of the role of Try, Ile or Leu in the action of Mel4 need to be confirmed in future studies that incorporate one or more of these amino acids in Mel4.

**Table 2.**
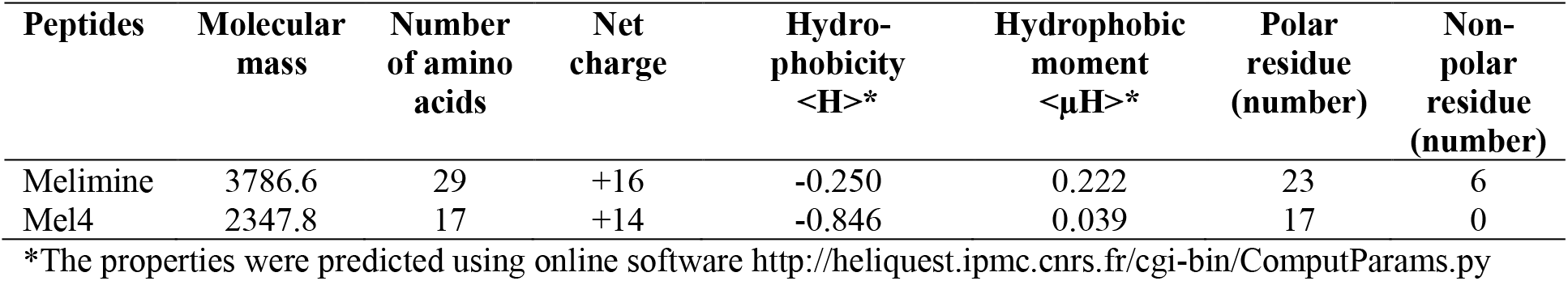
Properties of melimine and Mel4 peptides.

The membrane depolarization caused by both melimine and Mel4 was a concentration dependent event. These results are in agreement with previous studies showing a concentration-dependent depolarizing effect of the helical peptide LL-37 on *P. aeruginosa,* and PMAP-36, GI24 and melittin on *E. coli* cell membranes (54, 55). To further confirm whether membrane depolarization was a lethal event for these peptides, viable count was performed. Both AMPs reduced bacterial numbers upon membrane depolarization. Higher concentration of AMPs resulted in higher degrees of depolarization which in turn lead to larger amounts of bacterial death. Interestingly, sodium azide depolarization of the cell membrane did not affect cell viability. This suggest that membrane depolarization by itself was not sufficient to cause death of *P. aeruginosa.*

The membrane potential is essential for bacterial replication and ATP generation (56). Dissipation of the membrane potential may increase membrane permeability resulting in loss of cytoplasmic contents such as ATP (57). In the present study more than 70% ATP was released by melimine and approximately 35% by Mel4 following 2 minutes exposure, and the amount of ATP released plateaued at after four minutes exposure to both peptides. The effect of melimine is similar to the AMP porcine cathelicidin’s PR-39 effect of *E. coli* which induced 80% ATP leakage (58). The leakage of ATP results in depletion of intracellular ATP which in turn can lead to bacterial death (56). Interestingly, over the course of the study (10 mins), the amount of ATP released from cells during exposure to Mel4 never reached the amount released following exposure to melimine and also appeared to plateau indicating saturation. ATP released by bacteria can be depleted in supernatants of cells, possibly by hydrolysis at their cell surfaces (59). The data from the current experiments may indicate that cells treated with Mel4 retain some capacity to degrade extracellular ATP.

The permeability of cytoplasmic membrane also resulted in Sytox green uptake into cells. However, this uptake was relatively slow, and did not reach the level obtained with the positive control Triton X-100. Indeed, after 30 minutes incubation the amount of fluorescence due to Sytox green entering cells was equivalent to the amount that had entered cells after 10 minutes exposure to Triton X-100 when incubated with melimine, and <5 minutes when incubated with Mel4. Furthermore, with either melimine or Mel4 it took approximately 5 minutes for any fluorescence due to Sytox green to be registered. The difference in the time between membrane depolarization and Sytox green entering cells may be due to Sytox green requiring relatively large pore sizes in bacterial cell membranes for its uptake and time to intercalate with DNA. Compared with Sytox green uptake, ATP release was faster, and this may be due to the different mechanisms of entry/exit through cellular membrane. For example, the pore size needed for ATP to penetrate through membranes is a pore size of 1.5 nm (60), which is smaller than that needed for Sytox green to enter into cells. It may also take a longer time of Sytox green to cross the bacterial membranes and intercalate with DNA (61, 62). Similar kinetics to the current study of Sytox green staining resulting from interactions of the AMPs melittin and LL-37 has been reported against other Gram negative bacteria such as *P. aeruginosa, E. coli* and *Salmonella typhimurium* (63).

To further assess whether the peptides could damage the bacterial cell membrane, PI stain was used as an indicator of cells with ruptured membranes. Flow cytometry analysis indicated that treatment of *P. aeruginosa* with the both AMPs enhanced uptake of PI, suggesting that the bacterial cell membrane was disrupted. More than 50 % cells stained with PI after exposure to melimine for 30 minutes at its MIC. However, Mel4 took longer (150 min) for a similar amount of the cells to stain with PI at its MIC (**Fig. 3**). The positive control Triton-X resulted in less PI positive cells at this time point suggesting different rates of membrane permeabilization. Therefore, the sequence of steps occurring at the membrane, appears to begin with depolarization (with ATP and initial DNA/RNA leakage) and followed by more significant membrane disruption resulting in Sytox green and PI influx.

Melimine caused a concentration-dependent release of DNA/RNA (260nm absorbing material) as early as in 2 min, which is consistent with the results obtained by Minahk *et al.,* (64) who demonstrated a concentration-dependent release of DNA/RNA from *Listeria monocytogenes* after treatment with the antimicrobial peptide Enterocin CRL35 at the concentrations equal to its MIC and >4X its MIC. However, Mel4 induced release of DNA/RNA was dose independent. There was also a stepped release of DNA/RNA, with an initial burst release 2 minutes after addition of melimine or Mel4, and then another release of DNA/RNA between 60-150 minutes after addition of the peptides. The amount of DNA/RNA release caused by Mel4 did not reach the level released after exposure to melimine even at longer incubation times. AMPs have been shown to permeabilize bacterial membranes by forming morphologically diverse pores in terms of diameter, lipid conformation surrounding the pores, life span and threshold of AMPs require to stabilize the pores (65), and perhaps melimine and Mel4 form morphologically diverse pores which affect release of large molecules such as DNA/RNA. The two-step process of DNA/RNA release may be due to damage and disintegration of DNA/RNA within the cell over the 150 minutes incubation with the peptides allowing smaller lengths of DNA to exit cells. This may occur during so-called bacterial apoptosis-like death, mediated via *recA,* such as occurs during antibiotic-induced bacterial death (66). Examining changes in *recA* and the size of the liberated DNA/RNA in future experiments may help understand this further. In conclusion this study has revealed a comprehensive timeline of the mode of actions of melimine and Mel4 against *P. aeruginosa* involved disruption of the cell membranes, efflux of its intracellular contents and lysis of bacteria. It is likely that the amphipathic characteristics of melimine allowed disruption of the cell membrane more rapidly than Mel4 which only had very low amphipathicity.

## ACKNOWLEDGEMENTS

This work was supported by Australian Research Council (ARC) discovery project funding scheme (project number DP160101664). Authors are grateful to Christopher Brownlee of the Biological Resources Imaging Lab (BRIL) at the University of New South Wales, Australia for helping in Flow Cytometry analysis

